# A remarkable case of conserved domain swapping in the COMMD family of proteins

**DOI:** 10.64898/2026.02.02.703414

**Authors:** Ryan J. Hall, Meihan Liu, Manuela K. Hospenthal, Dion J. Celligoi, Rajesh Ghai, Kai-En Chen, Joanna Sacharz, David A. Stroud, J. Shaun Lott, Michael D. Healy, Brett M. Collins

**Author notes:** Corresponding authors: Michael D. Healy, Brett M. Collins,. These authors contributed equally (Ryan J. Hall, Meihan Liu).

## Abstract

The eukaryotic COMMD family of proteins are core subunits of the Commander protein complex, with a central role in endosomal membrane trafficking and signalling. Previous crystal and cryoEM structures show that COMMD and COMMD-like proteins form homo-oligomeric and hetero-decameric assemblies with a central ring structure formed by their small C-terminal COMM domains. Their α-helical N-terminal (HN) domains decorate each side of these rings, and in the eukaryotic Commander complex engage the coiled-coil proteins CCDC22 and CCDC93. Here, we have determined new crystal structures of the isolated HN domains of human Commd4, Commd9 and Commd10, and find that all three proteins form domain swapped structures with a remarkably consistent topology. This occurs via a conformational change in a hinge-loop between helices α2 and α3, leading to exchange of helices α3 to α6 between protomers. The hinge-loops of Commd9 and Commd10 possess several serine and threonine residues that can be phosphorylated, and we find that specific phospho-mimicking mutations in Commd10 can promote or inhibit domain swapping. Whether the unique COMMD HN domains play any roles beyond assembly with CCDC proteins is unclear, but this work suggests a common conformational switch exists with a potential to regulate their function.

## INTRODUCTION

Endosomes are critical cellular sorting stations in eukaryotes, regulating the transport of transmembrane proteins and lipids to degradative lysosomal compartments or mediating recycling of proteins to various destinations such as the Golgi and the plasma membrane^1–3^. The Commander complex is widely conserved among single and multicellular organisms and plays a prominent role in endosomal recycling to the cell surface. Commander is composed of sixteen subunits that are assembled in two sub-assemblies^4–8^. The Retriever sub-complex is structurally related to Retromer and is a trimer of VPS35L, VPS26C and VPS29^1^. The CCC sub-complex possesses two coiled-coil domain proteins CCDC22 and CCDC93, DENND10 and ten related subunits of the COMMD protein family. The COMMD (copper metabolism gene MuRR1 domain) proteins form the core of the CCC sub-complex, assembling into a heterodecameric ring structure that is intimately bound to the N-terminal regions of the CCDC22-CCDC93 dimer.

The ten COMMD proteins Commd1-10 all possess two small structural domains, an N-terminal helical (HN) domain and a C-terminal copper metabolism gene MuRR1 (COMM) domain, although the HN domain of Commd6 is missing in some species including humans^1,9–11^. The COMM domain forms an obligate dimeric structure, that then forms the core of a heterodecameric ring assembly^4–6,12^. The ten HN domains alternate around the periphery of the ring, and within the larger CCC complex primarily mediate interactions with N-terminal regions of the CCDC proteins. Recently it was shown that bacterial and archaeal species also possess single COMMD-like proteins with little sequence homology and an ability to form homo-oligomeric ring-shaped structures with the same overall architecture^13^. Mutations in subunits of the Commander complex, including the Commd4 and Commd9 proteins, lead to a congenital syndrome known as Ritscher-Schinzel syndrome (RSS) with cerebellar, cardiac, and craniofacial malformations and other phenotypes associated with liver, skeletal, and kidney dysfunction^14^. At the cellular level, Commander (and the COMMD subunits) is localised to actin-enriched endosomal domains^7,15–18^ where it is essential for recycling of various transmembrane proteins including lipoprotein receptors and integrins, recruited by the adaptor protein sorting nexin 17 (SNX17)^7,19–23^. Apart from their importance for Commander assembly and function, and potential novel roles in bacterial and archaeal species, the precise functions of the COMMD proteins are unclear.

The structures of human, archaeal and bacterial COMMD proteins are now relatively well understood from crystallographic, cryoelectron microscopy (cryoEM) and AlphaFold studies^4–6,12,13^. However, while studying the isolated HN domains of the human COMMD proteins we have uncovered a capacity for domain swapping that is remarkably similar for at least three of the family members, Commd4, Commd9 and Commd10. The domain swapping occurs via the same hinge-loop region between the second and third α-helices. The conformational changes induced by the domain swapping are generally incompatible with the COMMD structures within the functionally assembled Commd1-Commd10 heterodecamer. However, their consistent topology across distinct paralogues suggests a family-wide structural flexibility that may be important for their function.

## RESULTS

### Structural context of the COMM domain of human COMMD proteins

**Fig. 1A** depicts the domain organisation of the COMMD proteins, and **Fig. 1B** shows example oligomeric structures of both human and bacterial COMMD and COMMD-like proteins. The ten human COMMD proteins form a hetero-decameric ring assembly^4–6^, while bacterial and archaeal proteins form homo-octameric and homo-decameric structures^13^. Each COMMD monomer consists of the C-terminal COMM domain which forms the central ring of the complexes, and the N-terminal HN domain which decorates each side of the structures. In the human CCC assembly the ten COMMD family members are intimately entwined with the N-terminal disordered regions of the CCDC22 and CCDC93 proteins (**Fig. 1C**). The sequences of the eukaryotic HN domains are relatively well conserved in orthologues across different species (**Fig. 1D**; **Fig. S1A-S1B**) but are poorly conserved across homologues such as the ten human family members (**Fig. S1C**). The HN domain itself consists of seven α-helices, where the α4 helix forms the ‘spine’ of the domain and the α6 and α7 helices generally form a contiguous structure separated by a short kink (**Fig. 1B-1C**).

**Figure 1.**
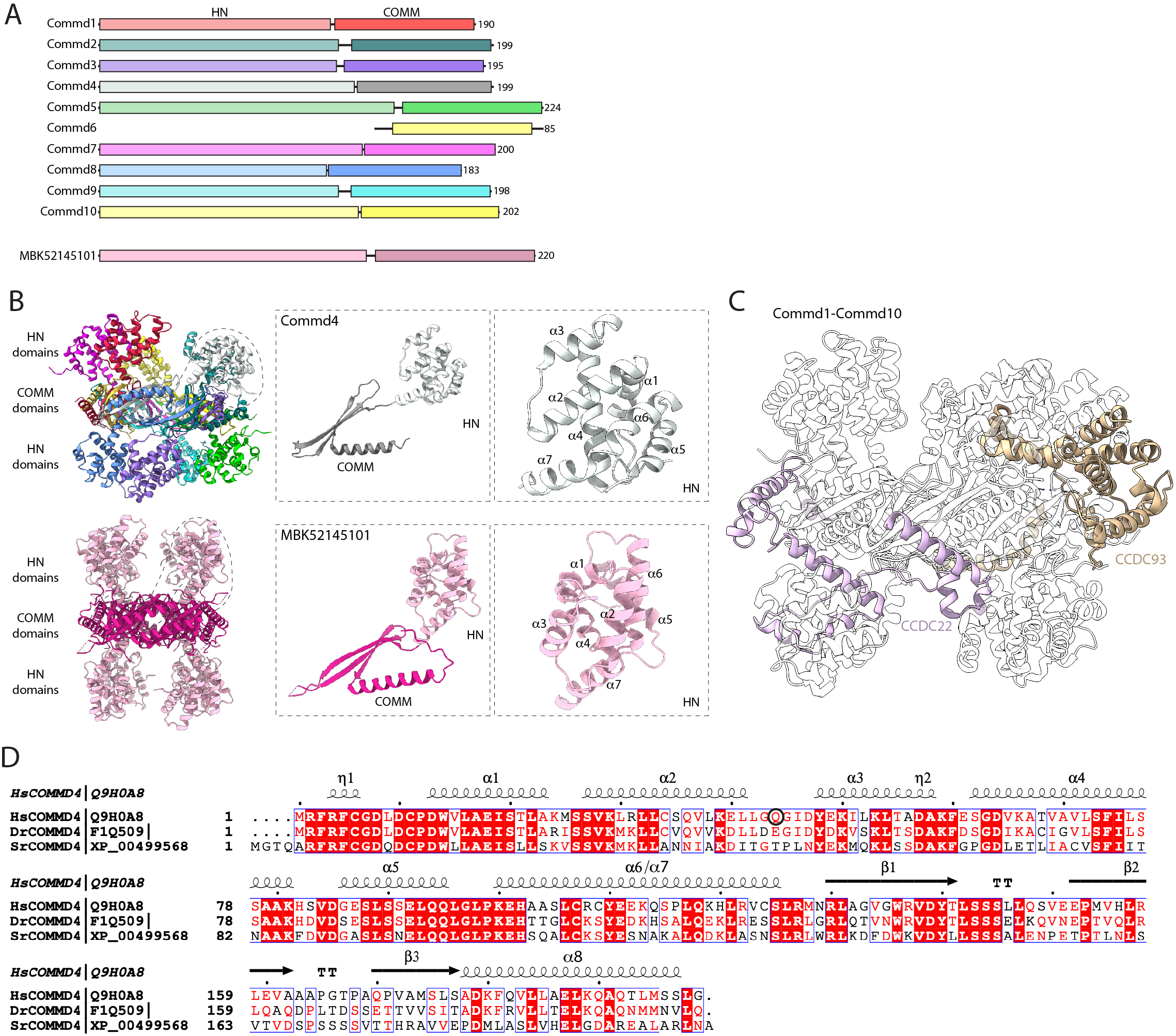
Structural context of the COMMD HN domains. **(A)** Schematic diagram showing to scale the domain organisation of the human COMMD family of proteins and the bacterial COMMD-like protein MBK52145101. Note that human Commd6 lacks the N-terminal HN domain although it is present in many other species. **(B)** Structures of the human Commd1-Commd10 heterodecameric complex (PDB ID 8F2R)^4,6^, and bacterial MBK52145101 homooctameric complex (PDB ID 9Q6C)^13^. The left panels show the overall assemblies, and the right panels show the structures of isolated monomers of Commd4 and MBK52145101 with enlarged views of their N-terminal HN domains. **(C)** Structure of the human Commd1-Commd10 heterodecameric complex with the N-terminal regions of the CCDC22 and CCDC93 proteins that form the CCC assembly. The COMMD proteins are shown in transparent ribbons to highlight the intimate entwining of the CCDC N-terminal regions. **(D)** Sequence alignment of Commd4 from humans, zebrafish and the unicellular choanoflagellate *Salpingoeca rosetta*. Secondary structure elements are shown for the human Commd4 protein.

### Commd4, Commd9, and Commd10 HN domains form domain swapped structures through a conserved mechanism

Although the HN domains of COMMD and COMMD-like proteins are not expected to function independently of the oligomerising COMM domains, we were interested to determine if they might adopt different conformations compared to what has been observed in previous crystal and cryoEM structures^4–6,13^. The nine HN domains of the human COMMD proteins were therefore cloned, expressed in bacteria and purified for biophysical and structural studies (**Fig. 2A; Table S1**). Although the HN domains could all be purified to homogeneity, several of the proteins also appeared to exist as larger oligomeric species. Specifically, while the isolated HN domains of Commd1, Commd2, Commd3, Commd5, Commd7 and Commd8 were primarily monomeric, the HN domains of Commd4, Commd9 and Commd10 existed in solution both as monomers and as larger species, although the amount of oligomeric Commd9 is substantially lower when compared to Commd4 and Commd10. Moreover, at least in the case of Commd4 these different species are relatively stable, as when each of two peaks is isolated and reanalysed (without being further concentrated) they remain in either monomeric or oligomeric peaks respectively (**Fig. S2**).

**Figure 2.**
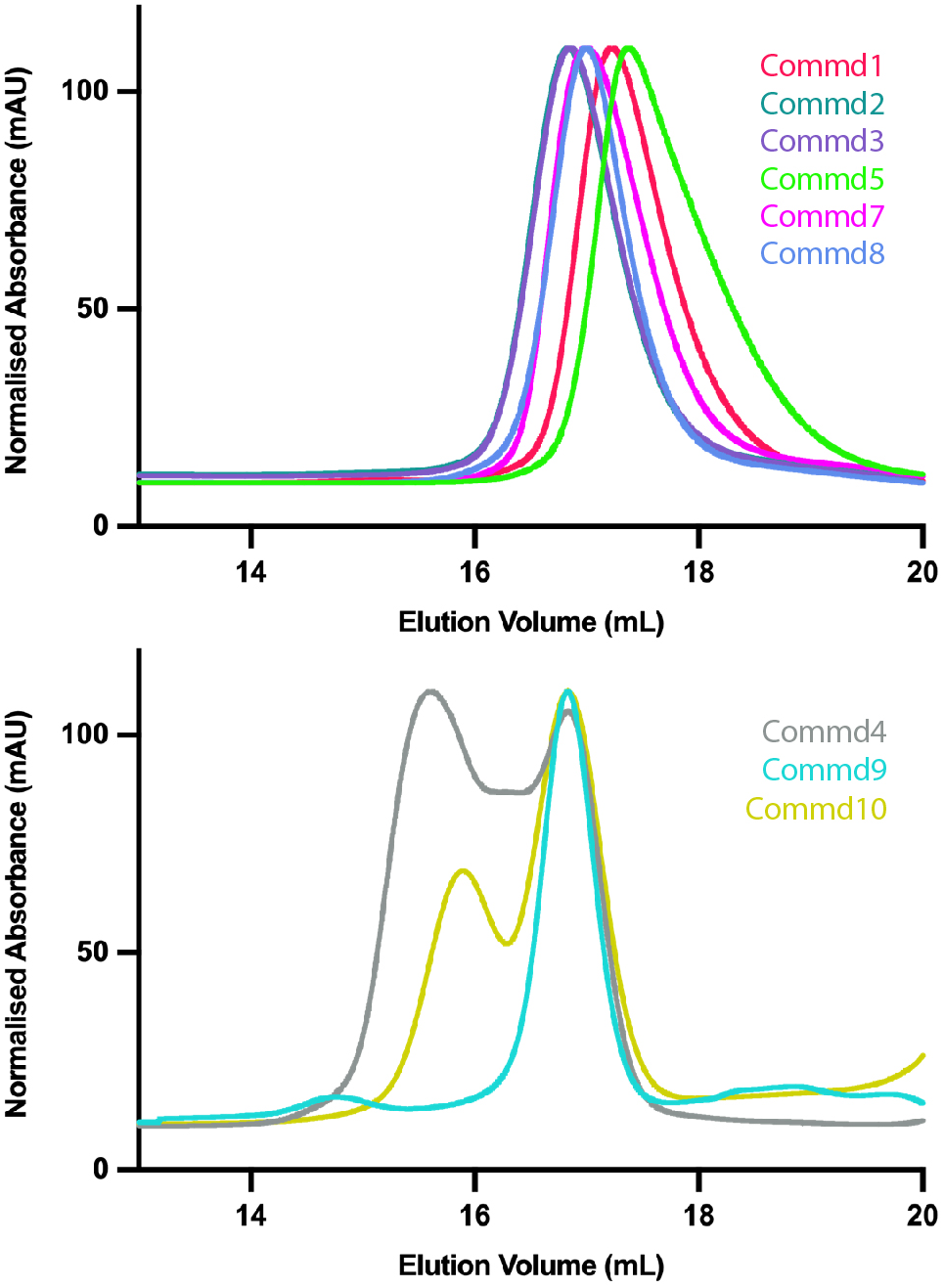
Analytical gel filtration the human COMMD HN domains. The HN domains of the nine human COMMD proteins were expressed in *E. coli*, and purified to homogeneity. Each protein was then analysed on a Superdex75 10/30 column to qualitatively examine their oligomeric states. The HN domains of Commd1, Commd2, Commd3, Commd5, Commd7 and Commd8 were predominantly monomeric in solution. The HN domains of Commd4, Comm9 and Commd10 displayed the presence of both monomeric and dimeric or trimeric oligomeric species.

To study how the oligomers of Commd4, Commd9 and Commd10 HN domains were assembled we determined their structures by X-ray crystallography (**Fig. 3A-3C**; **Table S2 and Table S3**). We were unable to solve the Commd10 HN structure by molecular replacement and therefore we used selenomethionine (SeMet) labelling and multi-wavelength anomalous diffraction (MAD) to determine its structure. Commd4 and Commd9 were both solved by molecular replacement, although Commd4 required multiple experiments with various fragment templates derived from the AlphaFold2 predicted model and previous cryoEM structures to eventually obtain a solution. In all three cases domain swapping has occurred, with clear electron density defining the swapped regions (**Fig. 3A-C; Fig. S3**). Previously we determined the crystal structure of the Commd9 HN domain in a monomeric state^12^, which closely matches its structure within the full-length protein studied by cryoEM^4–6^ (**Fig 3B**). However, here we report an alternate crystal form in which the asymmetric unit contained six copies of the Commd9 HN domain arranged in two trimeric domain swapped conformations.

**Figure 3.**
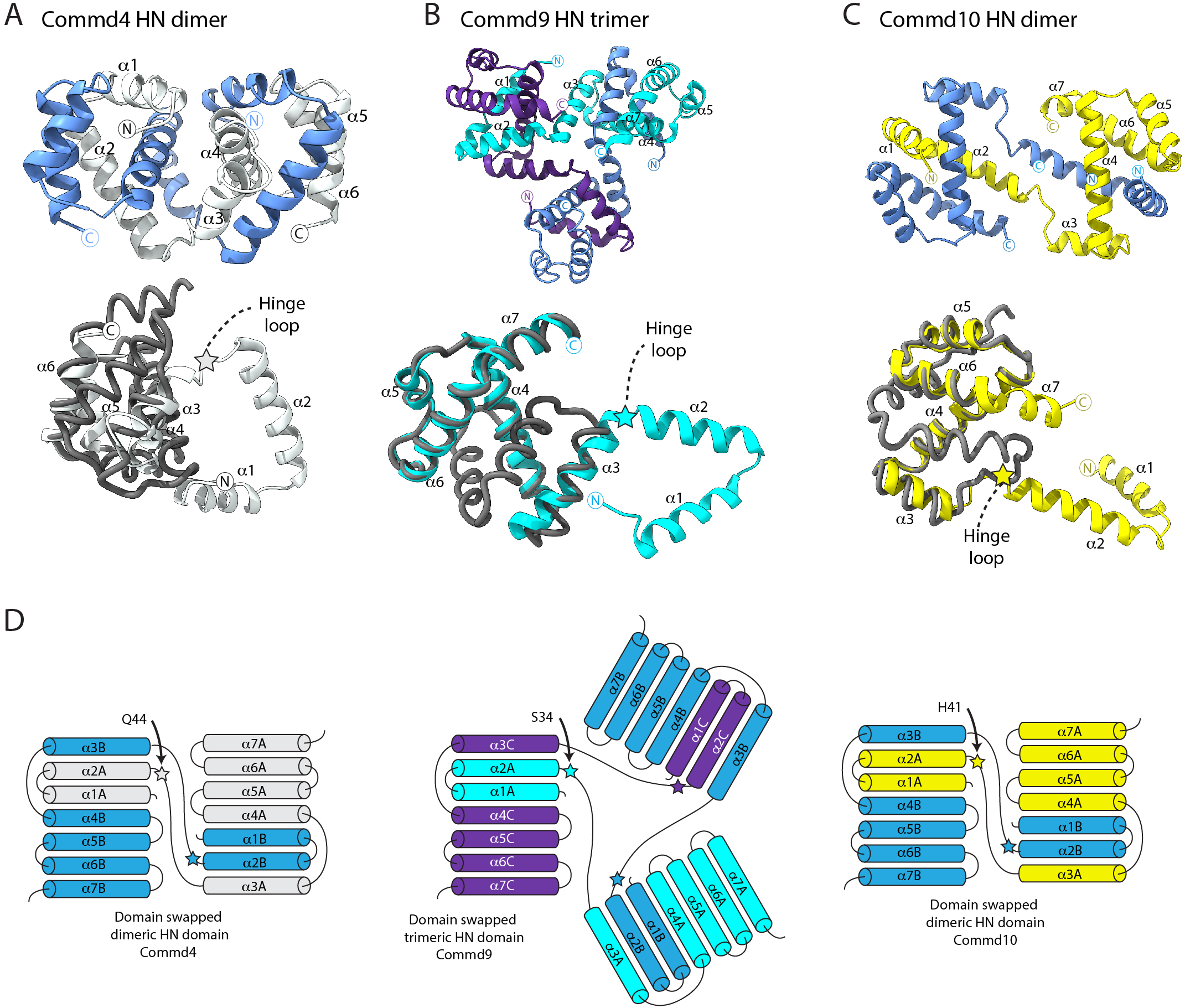
Structures of the Commd4, Commd9 and Commd10 domain swapped HN domains. **(A)** Crystal structure of the Commd4 HN domain showing the two domain swapped protomers in blue and white (top panel). The lower panel shows the domain swapped structure (white cartoon) aligned with the monomeric Commd4 HN domain (grey ribbon) derived from the Commd1-Commd10 heterodecameric complex (PDB ID 8F2R)^4,6^. **(B)** Crystal structure of the Commd9 HN domain showing the three domain swapped protomers in cyan, purple and blue (top panel). The lower panel shows the domain swapped structure (cyan cartoon) aligned with the monomeric Commd9 HN domain (grey ribbon) derived from the Commd1-Commd10 heterodecameric complex (PDB ID 8F2R)^4,6^. **(C)** Crystal structure of the Commd10 HN domain showing the two domain swapped protomers in cyan, purple and blue (top panel). The lower panel shows the domain swapped structure (cyan cartoon) aligned with the monomeric Commd10 HN domain (grey ribbon) derived from the Commd1-Commd10 heterodecameric complex (PDB ID 8F2R)^4,6^. **(D)** Topology diagrams of the domain-swapped oligomers of Commd4, Commd9 and Commd10 HN domains.

As shown in **Fig. 3A-C**, the domain swap in Commd4, Commd9 and Commd10 occurs via a remarkably similar mechanism. Although the precise organisation and number of subunits vary for the three proteins, the swap always occurs via a hinge-loop between the second and third α-helices shown schematically in **Fig. 3D**. This hinge-loop region, although it is in the same location, does not appear to possess a conserved sequence or structural motif across the three proteins (**Fig. S2C**). In Commd4 the structural divergence begins at Gln44, in Commd9 at Ser34 and in Commd10 at His41. If the α3-α7 helices are considered as one rigid body and the α1-α2 helices are treated as another, the overall structural transition can be described as involving a pivot or hinge-point between α2 and α3 from which the α1-α2 helices swing away (**Movies S1-S3**). These helices can then engage the α3-α7 helices of a second HN domain protomer. In the case of Commd4 and Commd10 there is a reciprocal exchange between two protomers to form a conjoined dimer. In the case of Commd9, the second protomer engages a third protomer which then closes the loop to form an interconnected trimer.

### Phosphorylation of the Commd10 hinge-loop can regulate its domains-swapping structure

Although the hinge-loop sites within Commd4, Commd9 and Commd10 do not share common sequences, we wondered if the conformational changes could be potentially regulated by a post-translational modification. According to the Phosphosite database (www.phosphosite.org) Commd4 can be ubiquitinated and/or acetylated at Lys39, Lys50 and Lys53 at the end of the α2 helix and Gln44 at the start of the α3 helix on either side of the hinge point. Similarly, Commd9 can be ubiquitinated at Lys40 adjacent to the hinge point at Ser34. In Commd10 Ser48 can be phosphorylated adjacent to the His41 hinge point, and both Commd9 and Commd10 possess multiple serines in the hinge-loop region. We decided to test if changing the hinge-loop serine residues in Commd10 to phosphomimetic glutamate residues would affect its capacity to dimerise *in vitro*. As shown in **Fig. 4A**, phosphomimetic mutations S47E and S50E both reduced the apparent dimerization of the Commd10 HN domain as the normal monomer/dimer mixed species observed by gel filtration now show a predominantly monomeric form of the protein. In contrast the S48E mutation shifts the ratio further towards the dimeric form (**Fig. 4A**), and when this peak is isolated and reanalysed, it remains stable as a dimer (not shown). We obtained crystals of each of the phosphomimic Commd10 HN domain mutants, but only crystals of the S50E mutant diffracted to high resolution. Although Commd10 HN(S50E) crystallised in a different space group, the refined structure revealed a homo-dimeric domain swapped protein identical to the wild-type HN domain (**Fig. 4B**). This shows that despite Commd10 HN(S50E) being primarily monomeric in solution it is still able to adopt a domain swapped configuration.

**Figure 4.**
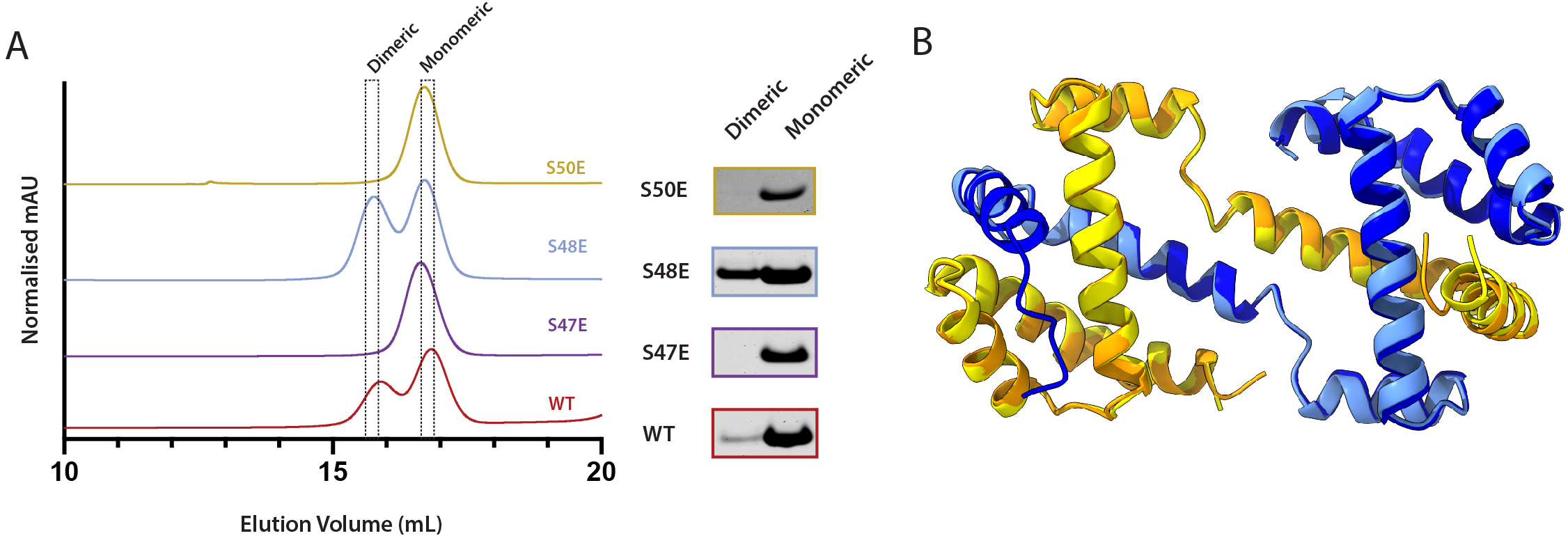
Phosphomimetic mutations in Commd10 inhibit but do not prevent domain swapping. (A) The wild-type and phosphomimetic mutants of Commd10 were analysed by gel filtration on a Superdex75 10/30 column to qualitatively examine their oligomeric states. The right panel shows Coomassie-stained gels of the two peak fractions for each sample. Mutations S47E and S50E reduce the amount of dimer formation, while the S48E mutant shows an increased proportion of dimeric species. **(B)** Crystal structure of Commd10 S50E HN domain (orange/dark blue) overlaid with the wild-type Comm10 HN domain (yellow/pale blue). Although the mutations inhibit dimer formation *in vitro*, it does not prevent domain swapping from occurring during crystallisation.

## DISCUSSION

This work has revealed an unexpected propensity for 3-dimensional (3D) domain swapping in the COMMD family of proteins. Conserved throughout evolution, and assembled into oligomeric rings^4–6,13^, the COMMD family have important roles in eukaryotic membrane trafficking and unknown functions in bacterial and archaeal species. Despite the known oligomeric assembly of COMMD proteins, 3D domain swapping has not previously been reported. The structures observed here show that the HN domains of Commd4, Commd9 and Commd10 (at least) are all able to exchange identical structural elements via a hinge-loop between the α2 and α3 helices, despite very little sequence similarity between them.

Domain swapping is a mechanism of dimer or oligomer formation that involves exchange of identical secondary or tertiary structural elements between protomers to form a stable inter-connected structure, where the topology of the original monomeric protein is preserved in the resulting structures. 3D domain swapping was first identified more than thirty years ago by the Eisenberg lab and occurs for a wide array of different folds^24,25^, with hundreds of examples available in the protein databank^26^. Although a frequently observed phenomenon, in most cases the functional importance of reported domain swapped structures is either unknown or remains speculative. Examples include the domain swapped interleukin-6 (IL-6) which may represent a functional dimeric state^27^, the homodimer of the pilotin InvH which may be displaced by the binding of the secretin InvG during formation of the Type III secretion system^28^, dimerisation of the DEP domain of Dishevelled thought to be important for Wnt signal transduction from the Frizzled receptor^29^, and evidence for a role of domain swapping in the evolution of the histone fold^30^. Uncontrolled domain swapping is also linked to protein misfolding and pathological protein aggregation including prions and amyloids^31–33^.

This raises the question – what is the functional significance of the 3D domain swapping observed for the human COMMD proteins? The major difficulty in studying the functional importance of domain swapping is that mutations designed to prevent domain swapping will also generally interfere with the folding of the monomeric protein. Another question that generally arises is whether a domain swapped structure can be regulated in some way, to control the conformational switching between monomeric and oligomeric species. One argument for a potential role of domain swapping in the COMMD proteins is that the same mechanism of fold switching occurs in three distinct homologues with largely varying sequences. But this is of course an argument based on structural correlations *in vitro* and there is currently no supporting evidence for these domain swapped structures *in vivo*. In addition, the domain swapped structures observed for the isolated HN domains would not easily be accommodated within the heterodecameric Commd1-Commd10 assembly without further conformational changes to avoid steric clashes or without compromising interactions with the CCDC22 and CCDC93 proteins (**Fig. S4**). Interestingly it was recently reported that COMMD3 might have Commander-independent function^34^, which raises the possibility of other Commander-independent roles where domain swapping may be of more importance. We speculated that post-translational modifications in the hinge-loop regions of the proteins could potentially regulate the domain swapped conformational changes, and showed that phosphorylation-mimicking mutations in Commd10 can either enhance or inhibit domain swapping from occurring in bacterially expressed proteins. The S48E mutation mimicking the confirmed phosphorylation site in Commd10 shifts the equilibrium towards the dimeric configuration, while S50E has the opposite effect. However, even though the Commd10 HN domain S50E mutant is predominantly monomeric in solution it still crystallised in the same domain swapped conformation suggesting these do not lead to all-or-nothing conformational changes.

In summary we have found that several members of the human COMMD protein family share a remarkable capacity for 3D domain swapping that occurs via the same mechanism but using distinct sequences. While the functional consequence of this fold-switching remains unclear, its conservation and potential regulation suggest an important structural mechanism for either creating conformational tension in COMMD subunits of the Commander complex, or regulating the assembly of this conserved family of proteins into oligomeric structures.

## MATERIALS AND METHODS

**Table S4** provides details of key resources.

### Molecular cloning

The bacterial expression constructs for the human COMMD HN domains were codon-optimised and synthesised by GeneUniversal (USA) and cloned into the pGEX6P-1 vector (Merck/Cytiva) at the BamHI and XhoI restriction sites. Bacterial and Archaeal COMMD HN domains were cloned into a modified pET28a vector with a GST tag using the golden gate protocol described by wicky *et al*^35^.

### Protein expression and purification

The bacterial expression plasmids encoding GST-tagged COMMD HN domains were transformed into Escherichia coli BL21-CodonPlus (DE3)-RIPL competent cells (Agilent). The bacterial cultures were grown in LB until OD600nm reached 0.6. The cultures were cooled to 18°C before inducing protein expression by adding 0.5 mM isopropylthio-β-galactoside (IPTG) and allowed to grow for 16 h. The cells were harvested by centrifugation at 6000 × g for 5 min at 4°C and the harvested cell pellet was resuspended in lysis buffer [20 mM Tris (pH 8.0), 500 mM NaCl, 10% glycerol, 50 μg/mL benzamidine, 100 units DNaseI, and 1 mM β-mercaptoethanol]. The cells were lysed by mechanical disruption at 27 kpsi using a Constant systems cell disrupter. The lysate was clarified by centrifugation at 50,000 × g for 30 min at 4°C. Proteins were purified using affinity chromatography from the clarified lysate.

GST-tagged proteins were purified on a glutathione-Sepharose (Cytiva) gravity column and the GST tag was cleaved with the addition of Prescission protease cleavage on to the beads with overnight incubation at room temperature. Finally, proteins were subjected to size exclusion chromatography using a Superdex 200 16/60 Hiload column or Superdex 75 16/60 Hiload column attached to an AKTA Pure (Cytiva) in 150 mM NaCl, 20 mM Tris (pH 8.0) and 1 mM dithiothreitol (DTT). The Commd9 HN domain with an N-terminal His-tag used for crystal structure determination was expressed in *E. coli* and purified as described previously^12^. The Commd10 HN domain was also transformed, expressed, grown in BL21-CodonPlus (DE3)-RIPL competent cells supplemented with seleno-L-methionine (SeMet) as reported previously^12^, using the method described by Van Duyne et al^36^. For analytical gel filtration chromatography proteins were run on a Superdex 200 10/30 column (Cytiva) in 150 mM NaCl, 20 mM Tris (pH 8.0) and 1 mM DTT. A similar SeMet protocol was used to express Commd4 HN however protein yield was not sufficient for crystallisation.

### Protein crystallisation and structure determination

The Commd4 HN domain was concentrated to 4 mg/ml and crystallised by vapour diffusion in 0.1 M Tris (pH 8.0) and 28% PEG4000. Data was collected at the Australian Synchrotron MX2 Beamline. Images were integrated with XDS^37^, and scaled and merged with AIMLESS^38^. The structure was determined by molecular replacement using PHASER^39^ with the Commd4 HN domain AlphaFold2 prediction^40,41^ as a search model. This required multiple searches using various fragments of Commd4 to obtain a solution producing reasonable electron density maps. The structure was refined with repeated rebuilding and refinement runs and model building with PHENIX^42^ and COOT^43^.

The Commd9 HN domain was concentrated to 12 mg/ml and crystallised by vapour diffusion in 20% PEG3350, 0.2 M Mg Nitrate (pH 8.0) at 18°C. Data was collected at the Australian Synchrotron MX2 Beamline. Images were integrated with XDS^37^, and scaled and merged with AIMLESS^38^. The structure was determined by molecular replacement using PHASER^39^ with the Commd9 monomeric HN domain (PDB ID 4OE9)^12^ as a search model. The structure was refined with repeated rebuilding and refinement runs and model building with PHENIX^42^ and COOT^43^.

The SeMet-labelled Commd10 HN domain was buffer-exchanged into 10 mM Tris (pH 8.0), 100 mM NaCl, 2 mM DTT, and concentrated to 20 mg/ml for crystallisation at 20°C. The protein was supplemented with 10 mM DTT before setting up hanging-drop crystallization screens using a mosquito liquid handling robot (TTP LabTech). Crystals of the HN domain were obtained in 30% PEG400 in 0.1 M Tris pH 8.0 in a hanging drop setup. Data was collected at the Australian Synchrotron MX2 Beamline. Images were integrated with XDS^37^, and scaled and merged with AIMLESS^38^. Datasets were examined for anomalous signal and any pathologies using Xtriage within the PHENIX suite^42^. The Commd10 HN domain was solved by multi-wavelength anomalous dispersion (MAD) using Autosol within the PHENIX software suite, including data collected at the peak, inflection and remote wavelengths for selenium atoms. The solution form Autosol was built using Autobuild and the resulting model was rebuilt with COOT^43^, followed by repeated refinement runs and model building with PHENIX^42^, ISOLDE^44^ and COOT^43^.

The Commd10 S50E HN domain mutant was concentrated to 14 mg/ml and crystallised by vapour diffusion in 30% PE14/4, 0.1 M magnesium formate, 0.1 M Tris (pH 8.5), with 10% glycerol added as cryopretectant. Data was collected at the Australian Synchrotron MX2 Beamline. Images were integrated with XDS^37^, and scaled and merged with AIMLESS^38^. The structure was determined by molecular replacement using PHASER^39^ with the Commd10 wild-type HN domain as a search model. The structure was refined with repeated rebuilding and refinement runs and model building with PHENIX^42^ and COOT^43^.

## Supporting information

Morph of Commd4 HN domain swap

Morph of Commd9 HN domain swap

Morph of Commd10 HN domain swap

## ACKNOWLEDGEMENTS

ray data were collected on the MX2 beamline at the Australian Synchrotron. The authors gratefully acknowledge the use of the Microscopy Australia Research Facility at the Centre for Microscopy and Microanalysis at The University of Queensland.

## AUTHOR CONTRIBUTIONS

Conceptualization, RJH, RG, JSL, BMC, MDH;

Methodology, ML, RJH, DJC, RG, JSL, BMC, MDH;

Investigation, YG, VAT, BMC, JR, YPW, TEH, MKH, DJC, JS;

Writing – Original Draft, ML, RJH, BMC, MDH;

Writing – Review & Editing, ML, RJH, JSL, BMC, MDH, DAS;

Funding Acquisition, JSL, BMC, DAS;

Supervision, JSL, RG, BMC, MDH.

## DATA AVAILABILITY

All data related to this study is available from the corresponding authors on request.

## Supplementary Information

**Figure S1.**
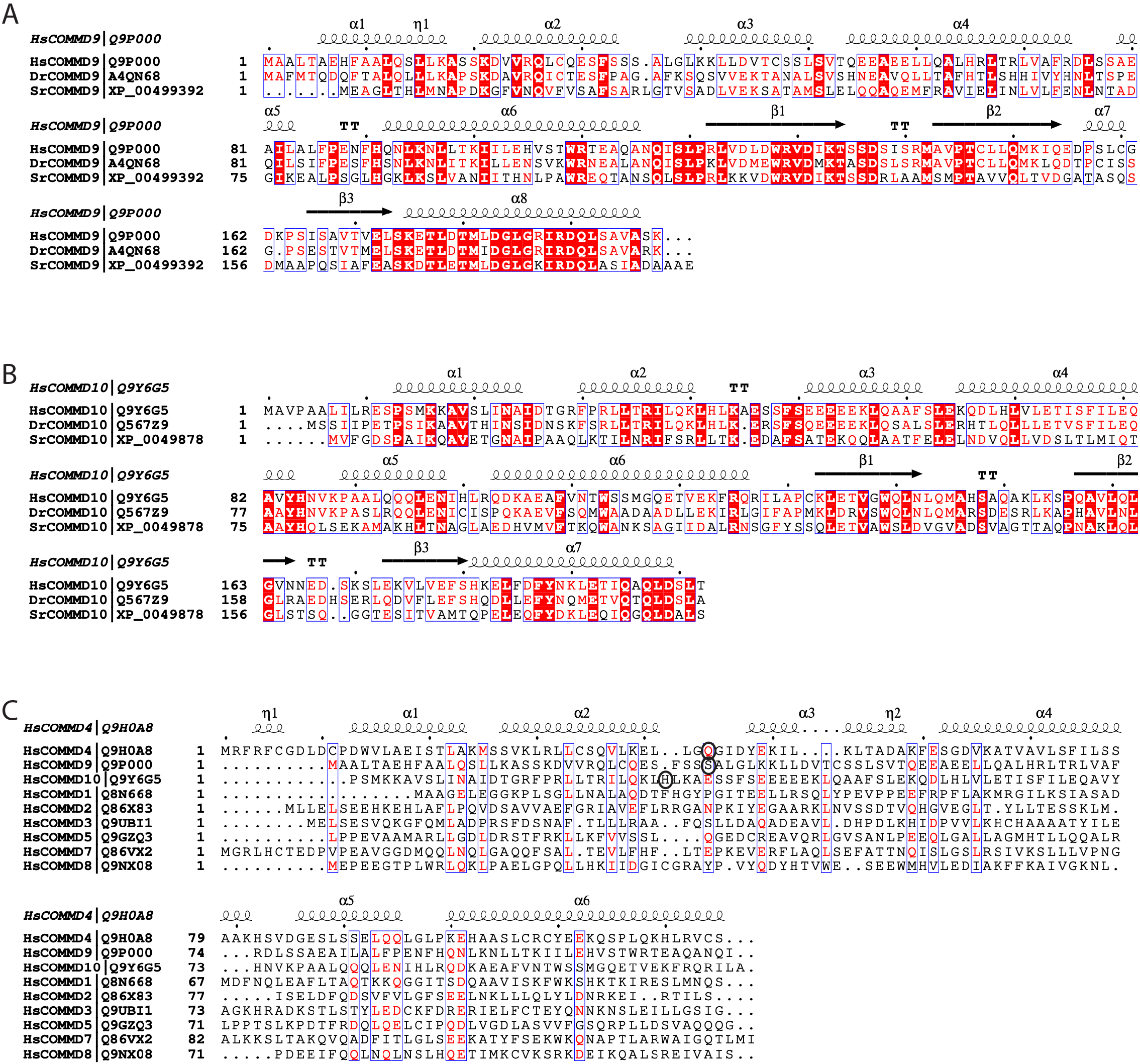
COMMD sequence alignments. **(A)** Sequence alignment of Commd9 from humans, zebrafish and the unicellular choanoflagellate *Salpingoeca rosetta*. Secondary structure elements are shown for the human Commd9 protein. **(B)** Sequence alignment of Commd10 from humans, zebrafish and the unicellular choanoflagellate *Salpingoeca rosetta*. Secondary structure elements are shown for the human Commd10 protein. **(C)** Sequence alignment of the human COMMD family of proteins. The secondary structure elements of Commd4 are shown at the top. Circles indicate the residues where the hinge-loop structural transition occurs for the domain swapping in Commd4, Comm9 and Commd10.

**Figure S2.**
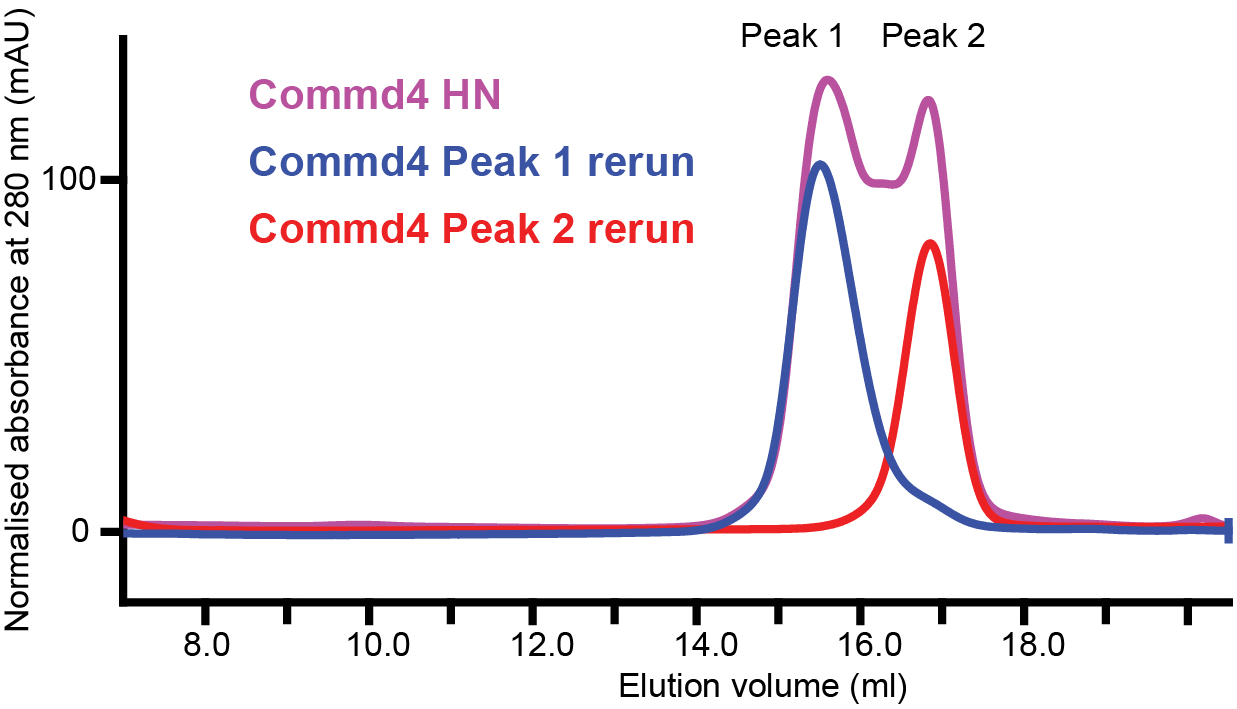
Analytical gel filtration of Commd4 HN domain. The HN domain of Commd4 was first isolated on a Superdex75 10/30 revealing both a monomeric (peak 2) and dimeric peak fractions (peak 1). When each peak fraction was rerun on the same column the protein remained stable in in their original monomeric or dimeric state.

**Figure S3.**
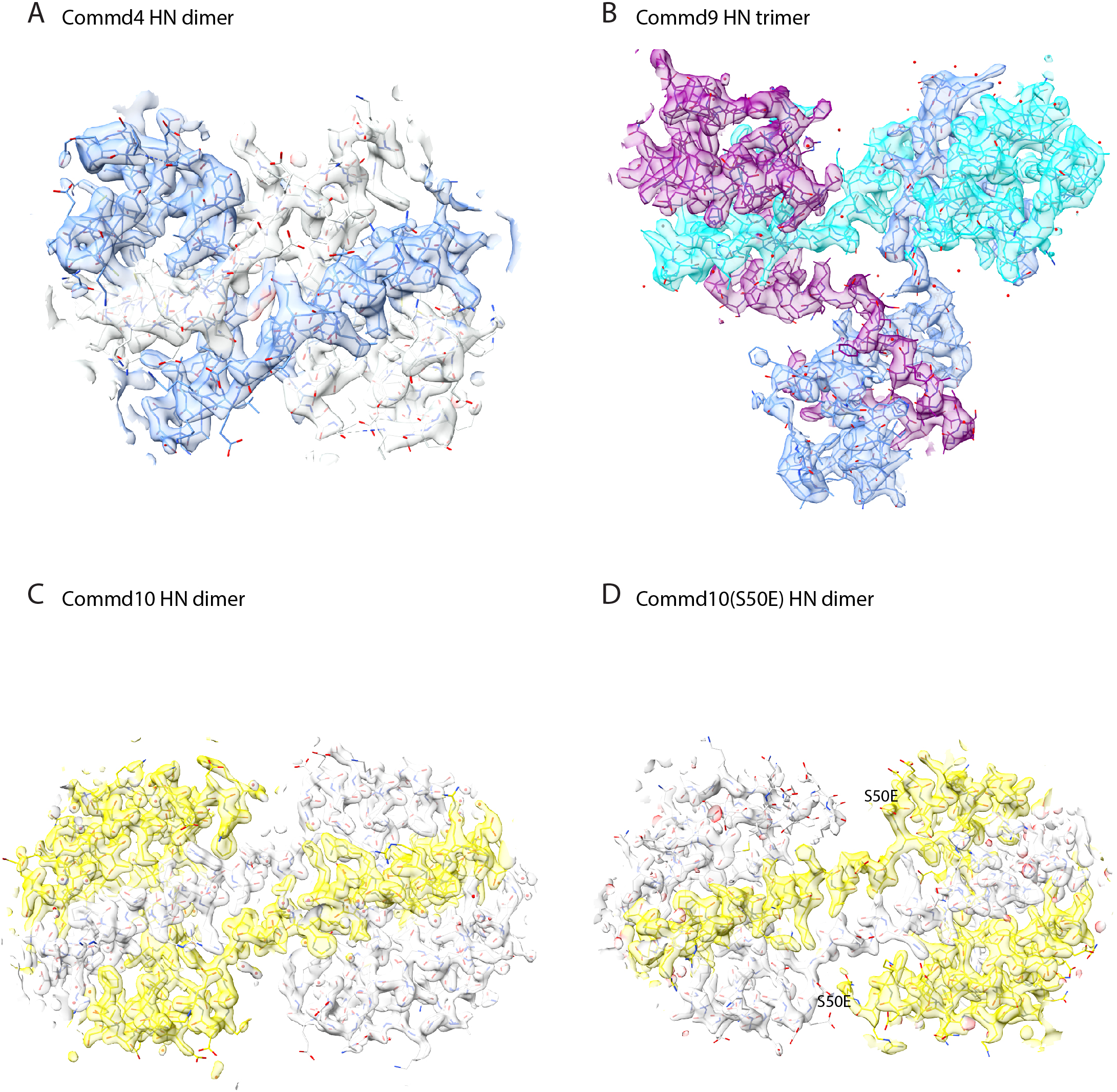
Electron density maps highlighting the domain swapped sequences of Commd4, Commd9 and Commd10 HN domains. Electron density maps for the HN domains of Commd4 **(A)**, Commd9 **(B)**, Commd10 **(C)** and Commd10 S50E mutant **(D)**. Figures show refined 2fo-fc maps contoured at 1.3σ. In each case the structures are oriented to highlight the extended regions where domain-swapping is occurring.

**Figure S4.**
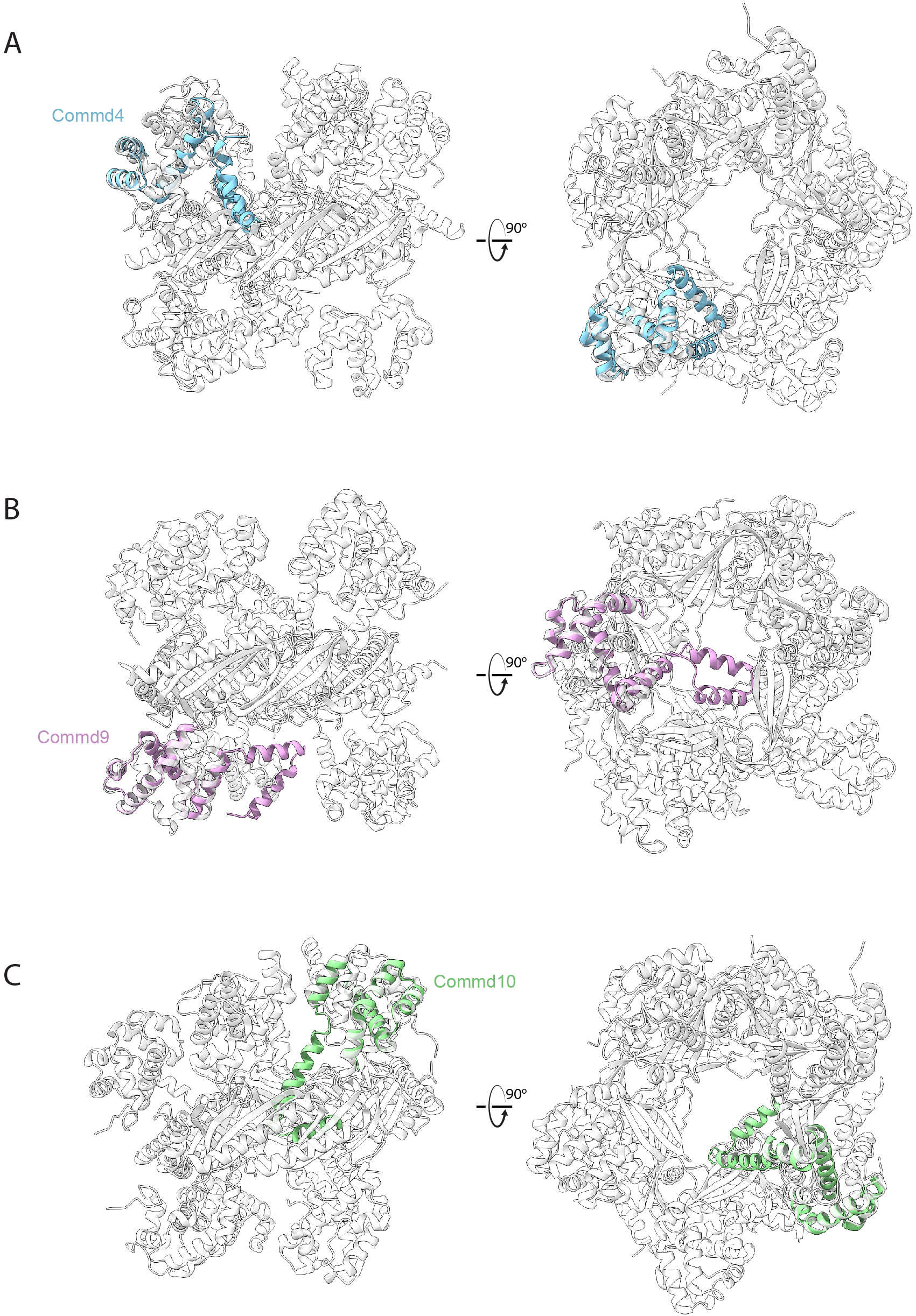
Structural alignments of the domain swapped HN domains. Structural alignments of the domain swapped HN domains of Commd4 **(A)**, Commd9 **(B)** and Commd10 **(C)** with the Commd1-Commd10 heterodecameric complex (PDB ID 8F2R)^4,6^. In each case the conformation of the domain swapped protomer would be sterically inconsistent with the overall assembly of the Commd1-Commd10 heterodecamer. In the case of Commd4 there would also be significant alterations in one of the major binding sites for the CCDC93 N-terminal domains.

**Table S1.**
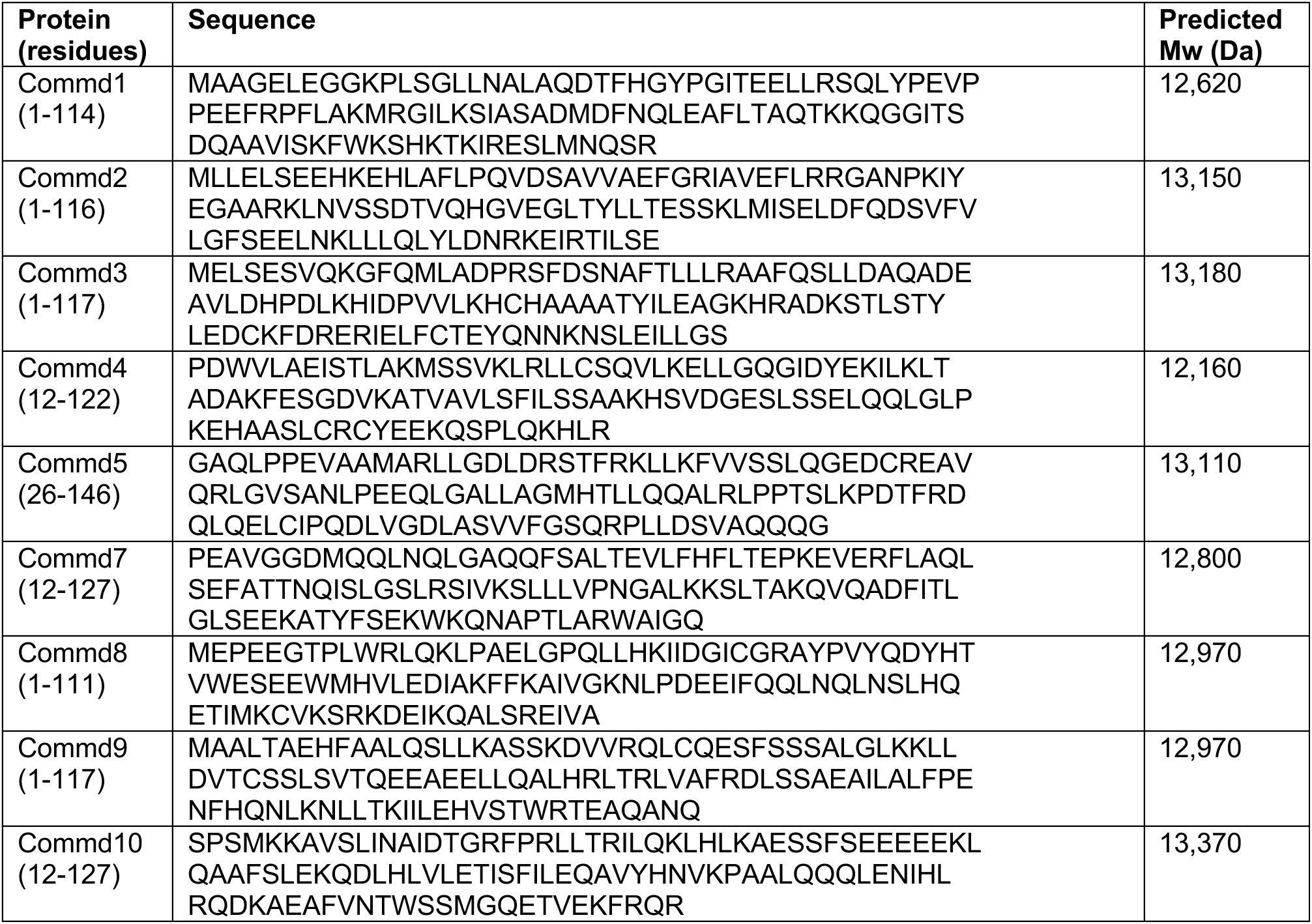
Sequences of the HN domains from human COMMD proteins examined.

**Table S2.** X-ray crystallographic structure determination of Commd10 HN domain.

|  | Commd10 HN domain<br>(peak) | Commd10 HN domain<br>(inflection) | Commd10 HN domain<br>(remote) | Commd10 HN<br>domain (native) |
| --- | --- | --- | --- | --- |
| **Data collection statistics** |  |  |  |  |
| PDB ID |  |  |  | 9ZMY |
| Space group | P2<sub>1</sub>2<sub>1</sub>2<sub>1</sub> | P2<sub>1</sub>2<sub>1</sub>2<sub>1</sub> | P2<sub>1</sub>2<sub>1</sub>2<sub>1</sub> | P2<sub>1</sub>2<sub>1</sub>2<sub>1</sub> |
| Unit cell dimensions<br>(a,b,c) (Å) | 40.8, 59.5, 96.8 | 40.8, 59.5, 96.8 | 40.8, 59.7, 96.7 | 41.6, 59.9, 96.9 |
| Unit cell angles (α,β,γ)<br>(°) | 90, 90, 90 | 90, 90, 90 | 90, 90, 90 | 90, 90, 90 |
| Wavelength (Å) | 0.97930 | 0.97960 | 0.96410 | 0.95372 |
| Resolution range (Å) | 40.7-2.80 (2.95-2.80) | 40.8-2.80 (2.95-2.80) | 48.4-2.80 (2.95-2.80) | 48.4-1.62 (1.64-1.62) |
| Total reflections | 87109 (12639) | 87129 (12604) | 87489 (12634) | 410343 (12613) |
| Unique reflections | 6220 (881) | 6223 (879) | 6244 (880) | 31360 (1141) |
| Multiplicity | 14.0 (14.3) | 14.0 (14.3) | 14.0 (14.4) | 13.1 (11.1) |
| Completeness (%) | 100.0 (100.0) | 100.0 (100.0) | 100.0 (100.0) | 98.8 (75.0) |
| Mean I/σ(I) | 16.0 (3.4) | 16.4 (3.5) | 7.9 (1.6) | 15.8 (1.1) |
| R-merge | 0.128 (0.852) | 0.123 (0.832) | 0.271 (2.225) | 0.051 (1.66) |
| R-meas | 0.137 (0.913) | 0.132 (0.892) | 0.290 (2.385) | 0.055 (1.81) |
| R-pim | 0.049 (0.329) | 0.035 (0.234) | 0.104 (0.329) | 0.015 (0.521) |
| CC(1/2) | 0.999 (0.929) | 0.999 (0.931) | 0.995 (0.858) | 1.00 (0.82) |
| R-work |  |  |  | 0.213 (0.367) |
| R-free |  |  |  | 0.241 (0.414) |
| Number of non-hydrogen atoms |  |  |  | 1926 |
| macromolecules |  |  |  | 1833 |
| water |  |  |  | 93 |
| RMS bonds (Å) |  |  |  | 0.006 |
| RMS angles (°) |  |  |  | 0.713 |
| Ramachandran favored<br>(%) |  |  |  | 99.1 |
| Ramachandran allowed<br>(%) |  |  |  | 0.9 |
| Ramachandran outliers<br>(%) |  |  |  | 0 |
| Clashscore |  |  |  | 1.9 |
| Average B-factor |  |  |  | 45.2 |

Values in parentheses refer to the highest resolution shell. $+R_{merge} = \sum |I - \langle I \rangle| / \sum \langle I \rangle$ , where $I$ is the intensity of each individual reflection. \* $R_{pim}$ indicates all $I^+$ & $I^-$ . $\frac{1}{2}R_{work} = \sum h |F_o - F_c| / \sum |F_o|$ , where $F_o$ and $F_c$ are the observed and calculated structure-factor amplitudes for each reflection $h$ . $R_{free}\%$ was determined by randomly selecting 10% of the diffraction data and excluded from refinement. Average $B$ ( $\text{\AA}^2$ )<sup>a</sup> was calculated using Baverage.

**Table S3.** Table S2. X-ray crystallographic structure determination of Commd4, Commd9 and Commd10(S50E) HN domains.

|  | Commd4 HN domain<br>(domain swapped dimer) | Commd9 HN domain (domain<br>swapped trimer) | Commd10 S50E HN domain<br>(domain swapped dimer) |
| --- | --- | --- | --- |
| **Data collection statistics** |  |  |  |
| PDB ID | 9ZMZ | 4NKN | 9ZN0 |
| Space group | P3<sub>1</sub>21 | I121 | P2<sub>1</sub> |
| Unit cell dimensions (a,b,c)<br>(Å) | 51.5, 51.5, 79.3 | 88.7, 79.8, 131.1 | 43.76, 92.17, 59.81 |
| Unit cell angles (α,β,γ) (°) | 90, 90, 120 | 90, 90.61, 90 | 90.00, 111.34, 90.00 |
| Wavelength (Å) |  | 0.95370 |  |
| Resolution range (Å) | 44.6-2.12 (2.18-2.12) | 32.79-2.79 (2.94-2.79) | 47.70-2.12 (2.18-2.12) |
| Total reflections | 139838 (9783) | 23054 (3158) | 95889 (7539) |
| Unique reflections | 7291 (562) | 6230 (853) | 24828 (1949) |
| Multiplicity | 19.1 (17.4) | 3.7 (3.7) | 3.8 (3.9) |
| Completeness (%) | 99.6 (95.8) | 98.6 (91.6) | 98.6 (95.1) |
| Mean I/σ(I) | 16.4 (1.6) | 16 (3.2) | 12.7 (3.7) |
| R-merge | 0.089 (1.51) | 0.093 (0.646) | 0.044 (0.263) |
| R-meas | 0.091 (1.558) | - | 0.051 (0.384) |
| R-pim | 0.021 (0.361) | - | 0.026 (0.152) |
| CC(1/2) | 0.999 (0.494) | - | 0.999 (0.950) |
| **Refinement statistics** |  |  |  |
| R-work | 23.0 (25.1) | 20.9 (31.6) | 19.4 (21.8) |
| R-free | 30.4 (37.1) | 24.6 (36.9) | 24.3 (28.2) |
| Number of non-hydrogen atoms |  |  |  |
| macromolecules | 751 | 4880 | 3721 |
| water | 13 | 61 | 125 |
| RMS bonds (Å) | 0.030 | 0.002 | 0.004 |
| RMS angles (°) | 2.52 | 0.396 | 0.858 |
| Ramachandran favored (%) | 96.8 | 99.8 | 98.2 |
| Ramachandran allowed (%) | 2.1 | 0.0 | 1.8 |
| Ramachandran outliers (%) | 1.1 | 0.2 | 0.0 |
| Clashscore | 5.2 | 3.0 | 3.6 |
| Average B-factor | 66.6 | 73.5 | 48.0 |

Values in parentheses refer to the highest resolution shell. $R_{merge} = \sum |I - \langle I \rangle| / \sum \langle I \rangle$ , where $I$ is the intensity of each individual reflection. $R_{pim}$ indicates all $I^+$ & $I^-$ . $R_{work} = \sum h |F_o - F_c| / \sum |F_o|$ , where $F_o$ and $F_c$ are the observed and calculated structure-factor amplitudes for each reflection $h$ . $R_{free}$ was determined by randomly selecting 10% of the diffraction data and excluded from refinement. Average $B$ ( $\text{\AA}^2$ )<sup>a</sup> was calculated using Baverage.

**Table S4.**
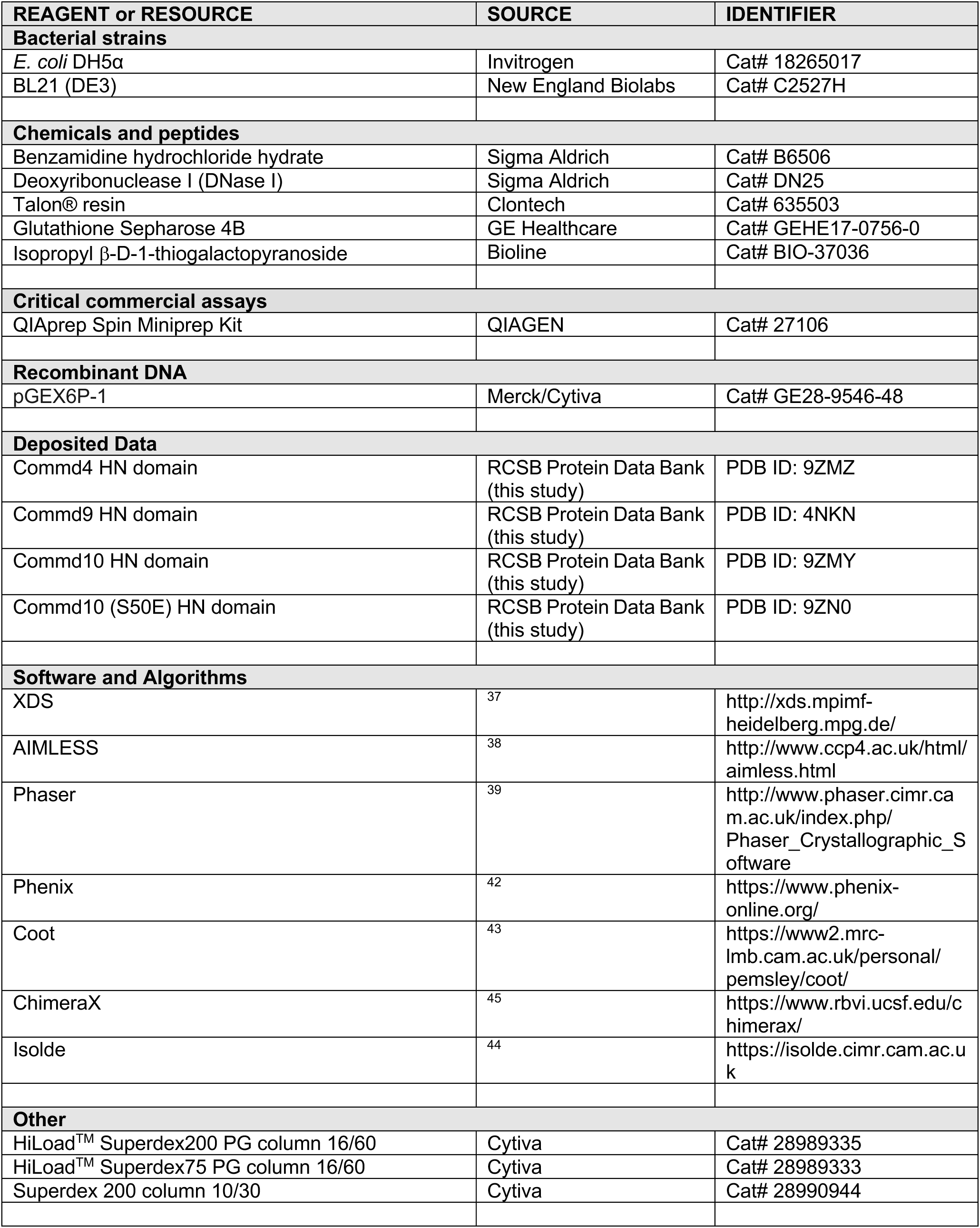
Key Resources.

